# Transcriptome analysis of a protein-truncating mutation in *sortilin-related receptor 1* associated with early-onset familial Alzheimer’s disease indicates effects on mitochondrial and ribosome function in young-adult zebrafish brains

**DOI:** 10.1101/2020.09.03.282277

**Authors:** Karissa Barthelson, Stephen Pederson, Morgan Newman, Michael Lardelli

## Abstract

The early cellular stresses which eventually lead to Alzheimer’s disease (AD) remain poorly understood because we cannot access living, asymptomatic human AD brains for detailed molecular analyses. Sortilin-related receptor 1 (*SORL1*) encodes a multi-domain receptor protein genetically associated with both rare, early-onset familial AD (EOfAD) and common, sporadic late-onset AD (LOAD). SORL1 has been shown to play a role in the trafficking of the amyloid β A4 precursor protein (APP) which is cleaved proteolytically to form one of the pathological hallmarks of AD, amyloid β (Aβ) peptide. However, the other functions of SORL1 are less well understood. Here, we employed a reverse genetics approach to characterise the effect of an EOfAD mutation in *SORL1* using zebrafish as a model organism. We performed targeted mutagenesis to generate an EOfAD-like mutation in the zebrafish orthologue of *SORL1*, and performed RNA-sequencing on mRNA isolated from a family of fish either heterozygous for the EOfAD-like mutation or their wild type siblings and identified subtle effects on the expression of genes which likely indicate changes in mitochondrial and ribosomal function. These changes appear to be independent of changes to expression of APP-related proteins in zebrafish, and mitochondrial content.

## Introduction

To reduce the prevalence and, therefore, the socioeconomic impacts of Alzheimer’s disease (AD), we must understand its molecular basis. Analysis of post mortem, human AD brains can give some insight into the disease mechanism. However, by the end stages of the disease, damage to the brain is considerable and little therapeutic intervention can be performed. An understanding of the early molecular changes, which occur decades before any symptoms arise, is necessary for development of effective treatments. We cannot access living, pre-symptomatic AD patient brain tissue for detailed molecular analysis. Therefore, analysis of animal models of AD is required.

A small number of genes are known to strongly influence the development of AD. Mutations in the presenilins (*PSEN1* and *PSEN2*) and the gene encoding amyloid β A4 precursor protein (*APP*) are causative for the early-onset (< 65 years of age), familial form of AD (EOfAD). EOfAD is an autosomal dominant disorder, and only accounts for a small proportion of all AD cases. The vast majority of AD cases arise sporadically, and have an age of onset later than 65 years (LOAD). Variation in at least 20 genes is associated with increased risk of developing LOAD, with the ε4 allele of the apolipoprotein E gene (*APOE*) contributing the greatest risk (1, 2). Since EOfAD and LOAD show similar disease progression and pathology (reviewed in (3)), analysis of EOfAD animal models may give insight into both of the main subtypes of AD. Intriguingly, sortilin-related receptor 1 (*SORL1*) appears to be associated with both EOfAD and LOAD (1, 2, 4-10) and may provide a mechanistic link between these two AD subtypes. However, *SORL1* is the least studied of the EOfAD genes and its precise role in AD is unclear.

The majority of research performed investigating the role of *SORL1* in AD is broadly based around its role in the trafficking of APP. It is well accepted that the binding of APP to SORL1 determines whether APP is directed through a recycling pathway, or is steered through the endolysosomal system to generate amyloid β (Aβ, the primary component of the senile neuritic plaques found in AD brains) (reviewed in (11)). SORL1 also binds Aβ itself, and guides it to the lysosome for degradation (12), and can act as a receptor for APOE (13). These findings have mostly been made using cell lines in which SORL1 has been overexpressed or removed. However, these manipulations do not closely reflect the pathophysiological state of *SORL1* in AD.

We recently submitted a paper investigating the effects on young adult zebrafish brain transcriptome state of heterozygosity for an EOfAD-like mutation and/or a putatively null mutation of the zebrafish orthologue of SORL1 (14). Heterozygosity for the EOfAD-like mutation resulted in subtle effects on the transcriptome, with only one gene detected as differentially expressed. At the pathway level, we found evidence for novel cellular processes previously unknown to require *sorl1*, such as energy metabolism, protein translation and degradation. These effects also were observed in the brains of fish heterozygous for a putatively null mutation in *sorl1*, suggesting they are due to haploinsufficiency. Transheterozygosity for the EOfAD-like and null mutations appeared to affect iron homeostasis and other cellular processes distinct from those detected in the heterozygous fish brains (14).

Here, we aimed to further our understanding of the effects of EOfAD mutations in *SORL1* by generating and analysing an additional zebrafish *sorl1* mutation model. We performed targeted mutagenesis to generate a line of zebrafish carrying the W1818* mutation in *sorl1*, which models the human *SORL1* EOfAD mutation W1821* (4). We then compared the brain transcriptomes of young adult *sorl1*^*W1818\**^ heterozygous and wild type sibling fish to identify, in an unbiased and objective manner, the changes in gene expression caused by the mutation. Consistent with our previous study, heterozygosity for the W1818* mutation resulted in subtle changes to brain gene expression. Genes involved in energy production and protein translation showed altered expression. The W1818* mutation of *sorl1* did not appear to affect Appa/Appb protein levels, cellular mitochondrial content, or iron homeostasis.

## Results

### Generation and characterisation of the EOfAD-like mutation W1818* in zebrafish *sorl1*

In 2012, Pottier et al. (4) characterised mutations in *SORL1* which segregate with EOAD in families with autosomal dominant inheritance patterns. We previously analysed the effect of a protein-truncating mutation from this study (human C1478*) on the brain transcriptome in a zebrafish model (zebrafish V1482Afs) (14). To further our understanding of the effects of protein-truncating mutations in *SORL1*, we generated a zebrafish model of an additional protein-truncating mutation, W1821* (4). The W1821 site is conserved in zebrafish (W1818). We edited this site in the zebrafish genome using targeted mutagenesis (**Additional File 1**) and isolated a line of zebrafish carrying an 11 nucleotide deletion, resulting in the equivalent protein level change observed in the human mutation (**Figure 1A)**.

**Figure 1:**
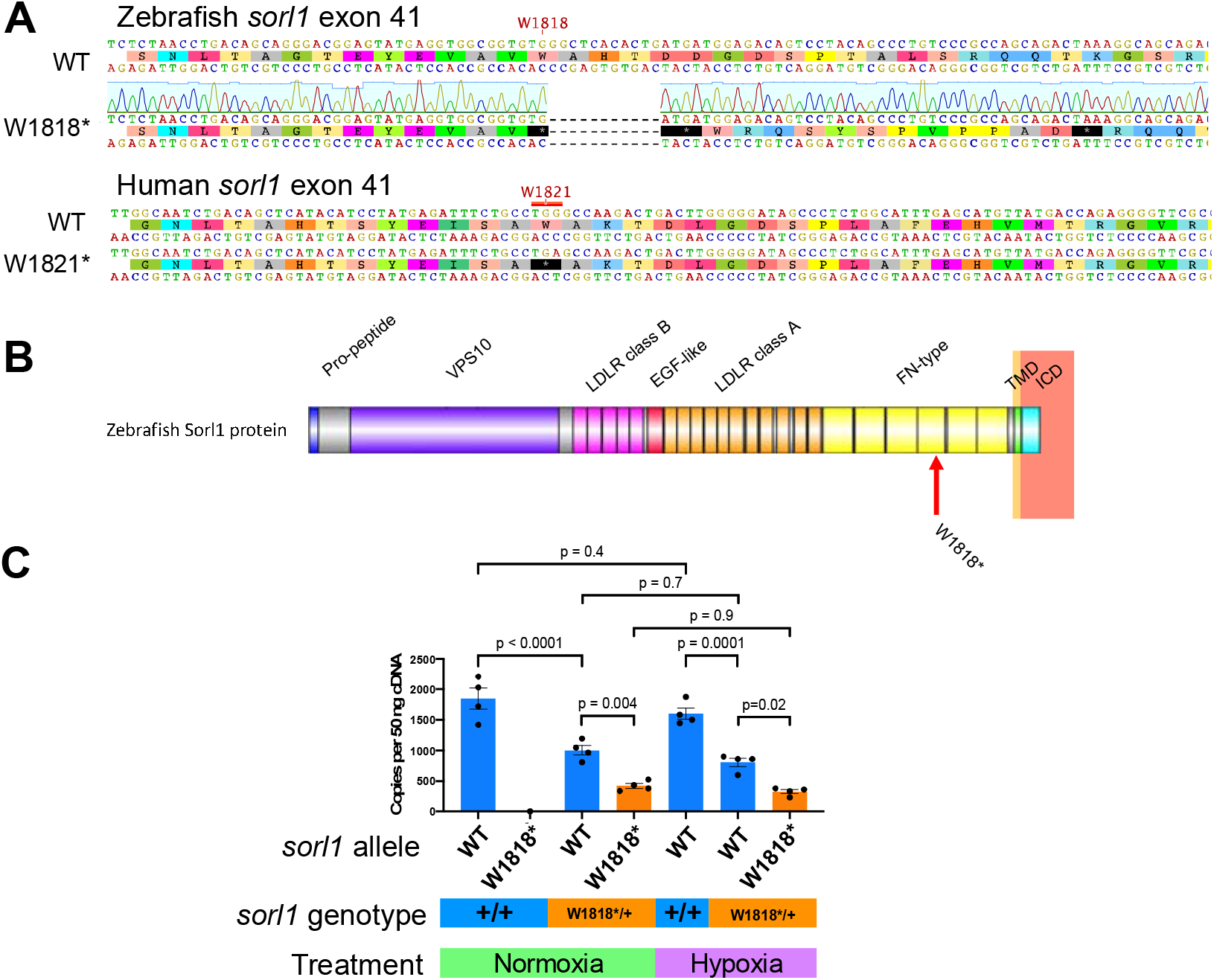
The W1818* mutation in *sorl1* results in a transcript subject to nonsense mediated mRNA decay. **A**. Alignment of zebrafish and human SORL1 wild type and mutant exon 41 sequences. **B**. Schematic of Sorl1 protein with protein domains and the site of the W1818* mutation indicated. VPS10: vacuolar protein sorting 10 domain. LDLR: low density lipoprotein receptor. EGF: epidermal growth factor. FN: fibronectin. TMD: transmembrane domain. ICD: intracellular domain. **C**. Number of copies of the *sorl1* wild type (WT) and mutant (W1818*) transcripts in 6 month old wild type (+/+) and heterozygous mutant (W1818*/+) sibling brains under normoxia and acute hypoxia. P-values were determined by two-way ANOVA with Tukey’s post hoc test for multiple comparisons.

Protein-truncating mutations in *SORL1* have been shown to be subject to nonsense mediated mRNA decay (NMD) in lymphoblasts from human carriers (4). Additionally, *SORL1* expression has been shown to be upregulated under hypoxia *in vitro* (15). Therefore, we asked whether the W1818* transcript of *sorl1* was also subject to NMD, and whether this was altered by hypoxia treatment. To address this, we performed allele-specific, digital quantitative PCRs (dqPCRs) on brain-derived cDNA from fish exposed to normoxia and acute hypoxia. We found that the W1818* transcript is less abundant in W1818*/+ brains relative to the wild type transcript under both normoxia (p < 0.0001) and acute hypoxia (p = 0.0001), supporting that the W1818* transcript is likely subject to NMD. We also observed that the wild type transcript in W1818*/+ brains is less abundant than in +/+ brains under normoxia (p = 0.004) and acute hypoxia (p = 0.02). We did not observe any significant change of *sorl1* transcript levels between normoxia and acute hypoxia (**Figure 1C**), despite that these fish were likely showing a transcriptional response to hypoxia, indicated by upregulation of *pdk1* (**Additional File 2**).

To determine whether the W1818* mutation alters the expression of Sorl1 protein, we performed western blot analysis on zebrafish brain lysates from +/+, W1818*/+ and W1818*/W1818* fish at 6 months of age. We immunoblotted with an antibody raised against the C-terminus of human SORL1 which cross-reacts with zebrafish Sorl1 protein. We observed loss of the signal observed at ∼ 250kDa in homozygous mutant brains, confirming that the W1818* mutation of *sorl1* results in disruption of translation of Sorl1 protein (**Additional File 3**).

In summary, *sorl1* does not appear to be upregulated by acute hypoxia *in vivo*, the W1818* mutation in *sorl1* results in a transcript which is likely subject to NMD, and the mutation disrupts wild type Sorl1 protein translation.

### Transcriptome analysis of W1818*/+ and +/+ young adult zebrafish brains

Which genes are dysregulated as a result of heterozygosity for the EOfAD-like, W1818* mutation in *sorl1* in young adult brains? To address this question, we performed mRNA sequencing on the brains from fish either heterozygous for the W1818* mutation and their wild type siblings at 6 months of age (n = 6).

The overall similarity between gene expression profiles can be explored using principal component analysis (PCA). Samples which have similar gene expression profiles will cluster together in a PCA plot. In our RNA-seq dataset, no clear separation of male and female samples was observed, supporting our previous observations that sex does not have a large effect on the zebrafish brain transcriptome (14). We observed partial separation of W1818*/+ and wild type samples across principal component 2 (PC2), accounting for approximately 16% of the total variation in this dataset (**Figure 2A**). Therefore, heterozygosity for the W1818* mutation likely does not have widespread effects on the transcriptome of young adult zebrafish brains.

**Figure 2:**
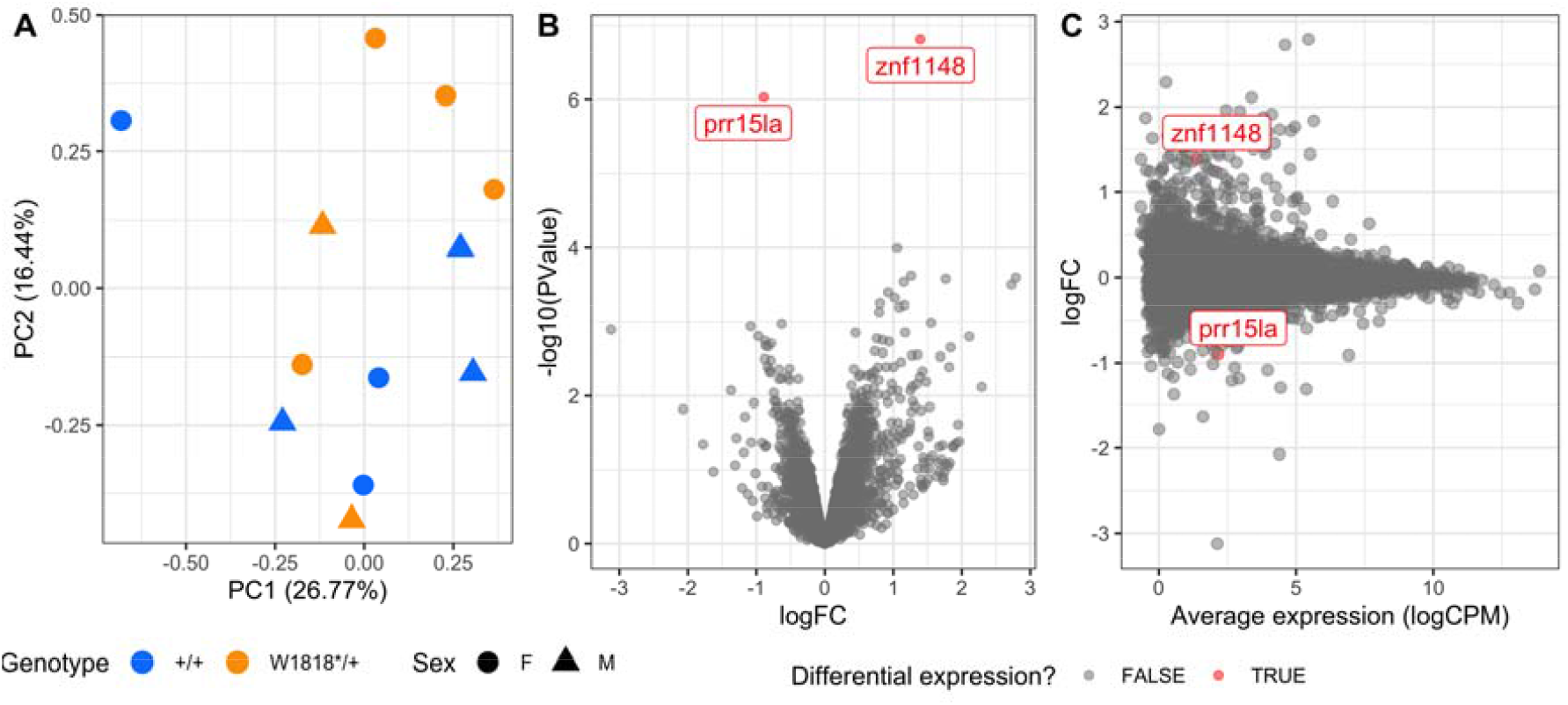
Heterozygosity for the W1818* mutation in *sorl1* has subtle effects on young-adult brain transcriptomes. **A**. Plot of principal component 1 (PC1) against PC2 from a principal component analysis. Circles indicate female samples (F) and triangles indicate male (M) samples. **B**. Volcano plot showing the significance (-log10(PValue)) and log_2_ fold change (logFC) of genes in W1818*/+ brains relative to +/+ sibling brains. The genes identified as significantly differentially expressed are coloured in red. **C**. Mean-difference (MD) plot of gene expression in W1818*/+ brains relative to +/+ sibling brains.

Differential gene expression analysis supported our conclusions from the PCA. Only two genes were identified as differentially expressed to a statistically significant degree due to *sorl1* genotype: zinc finger protein 1148 (*znf1148*) and proline rich 15 like a (*prr15la*) (**Figure 2B, 2C, Additional File 4**). The functions of these genes are not known. BLAST searches identified *ZNF99* and *PRR15L* as candidate human orthologues of *znf1148* and *prr15la* respectively. However, broad-scale analysis of conservation of synteny between the genomic regions occupied by the zebrafish and human genes was unable to support these orthologies (**Additional File 5**).

To obtain a more complete view of changes to brain gene expression due to heterozygosity for the W1818* mutation in *sorl1* we performed enrichment testing on particular gene sets (below) using the entire list of detectable genes in the RNA-seq experiment. For insight into the cellular processes represented by the gene expression changes we used the *KEGG* and *HALLMARK* gene sets from the *Molecular Signatures Database* (*MSigDB*, (16, 17)). To test for possible iron dyshomeostasis, we used our recently defined gene sets representing the genes containing iron-responsive elements (IRE) in the untranslated regions (UTRs) of their mRNAs (18). We applied the self-contained gene set testing methods *fry* (19) and *GSEA* (17, 20), and the competitive gene set testing method *camera* (21), and combined the resulting p-values by calculating the harmonic mean-p value, a recently developed method of combining dependent p-values (22). We further protected from type I errors by performing an FDR adjustment on the resulting harmonic mean p-value.

In all, we identified six gene sets significantly altered as a group after FDR adjustment of the harmonic mean p-value (**Figure 3**). However, the leading edge genes from the *GSEA* algorithm (which can be thought of as those genes driving the enrichment of the gene set) for four of these six gene sets (*HALLMARK_* & *KEGG_OXIDATIVE_PHOSPHORYLATION, KEGG_HUNTINGTONS_DISEASE*, and *KEGG_PARKINSONS_DISEASE*) all share many genes, showing that the statistical significance of these gene sets is, essentially, being driven by the same signal (genes encoding components of the mitochondrial electron transport chain). No IRE gene sets were found to be significantly altered, consistent with our previous observations for heterozygosity for an EOfAD-like allele of *sorl1* (14).

**Figure 3:**
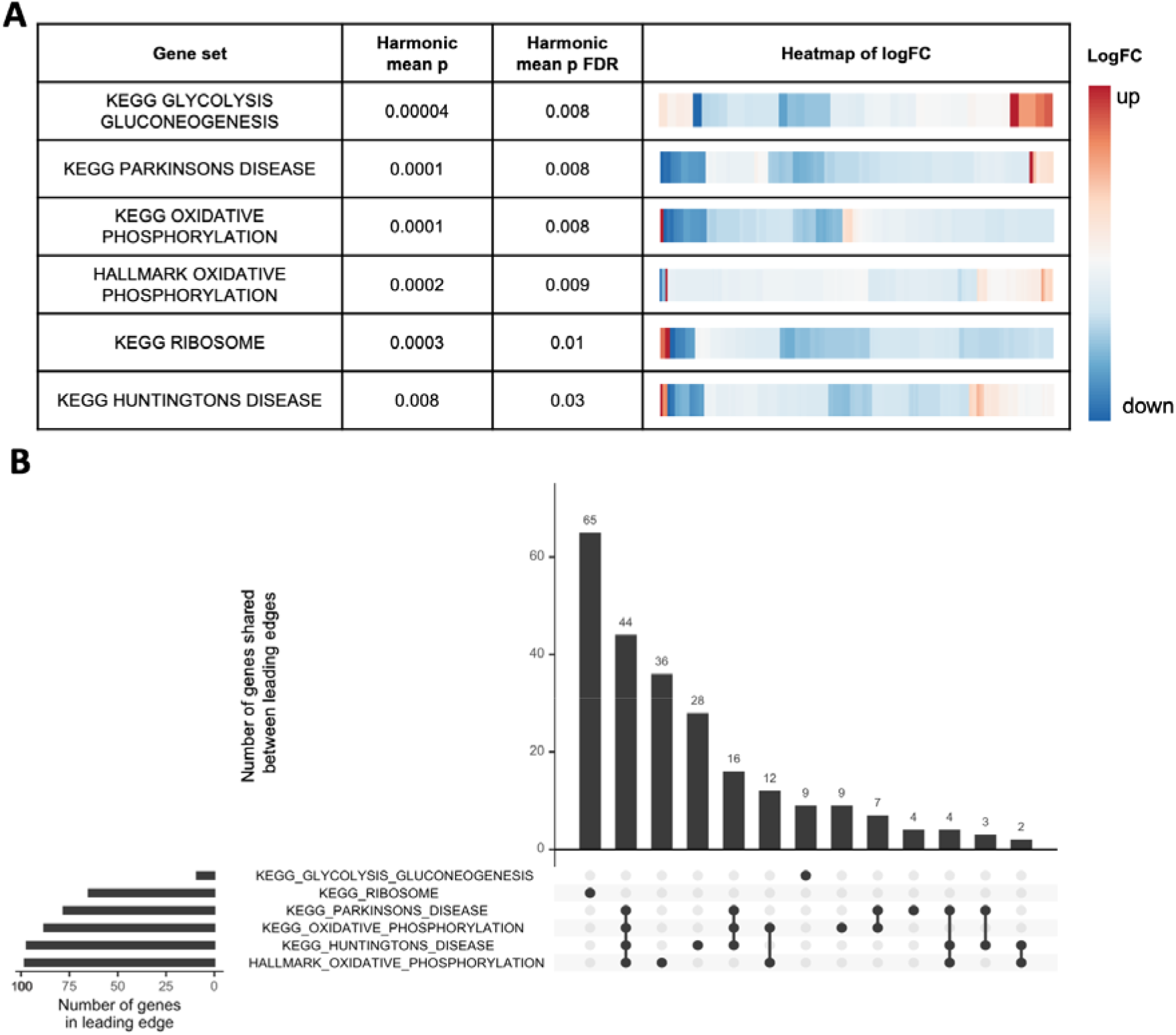
Mitochondrial and ribosomal function are predicted to be affected by heterozygosity for the W1818* mutation in *sorl1*. **A**. Table indicating the significantly altered KEGG and HALLMARK gene sets (FDR adjusted harmonic mean p-value < 0.05) in W1818*/+ brains relative to +/+ brains. Heatmaps indicate the log2 fold change (logFC) of all detectable genes within the gene sets, clustered by their Euclidean distance. **B**. Upset plot of the leading edge genes from the GSEA algorithm for the significant gene sets indicating the number of shared genes driving the enrichment of the gene set.

Since our transcriptome analysis was of bulk mRNA isolated from entire zebrafish brains, differences in cell type proportions between W1818*/+ and +/+ brains could cause artefactual changes in gene expression levels. To investigate this possibility, we visualised the expression levels of marker genes present in four broad cell types within zebrafish brains: neurons, astrocytes, oligodendrocytes (23) and microglia (24) (**Figure 4A**). Since we did not observe any obvious differences in marker gene expression levels between genotypes, it is unlikely that changes in cell type proportions cause the differential expression of genes observed in the W1818* heterozygous fish brains.

**Figure 4:**
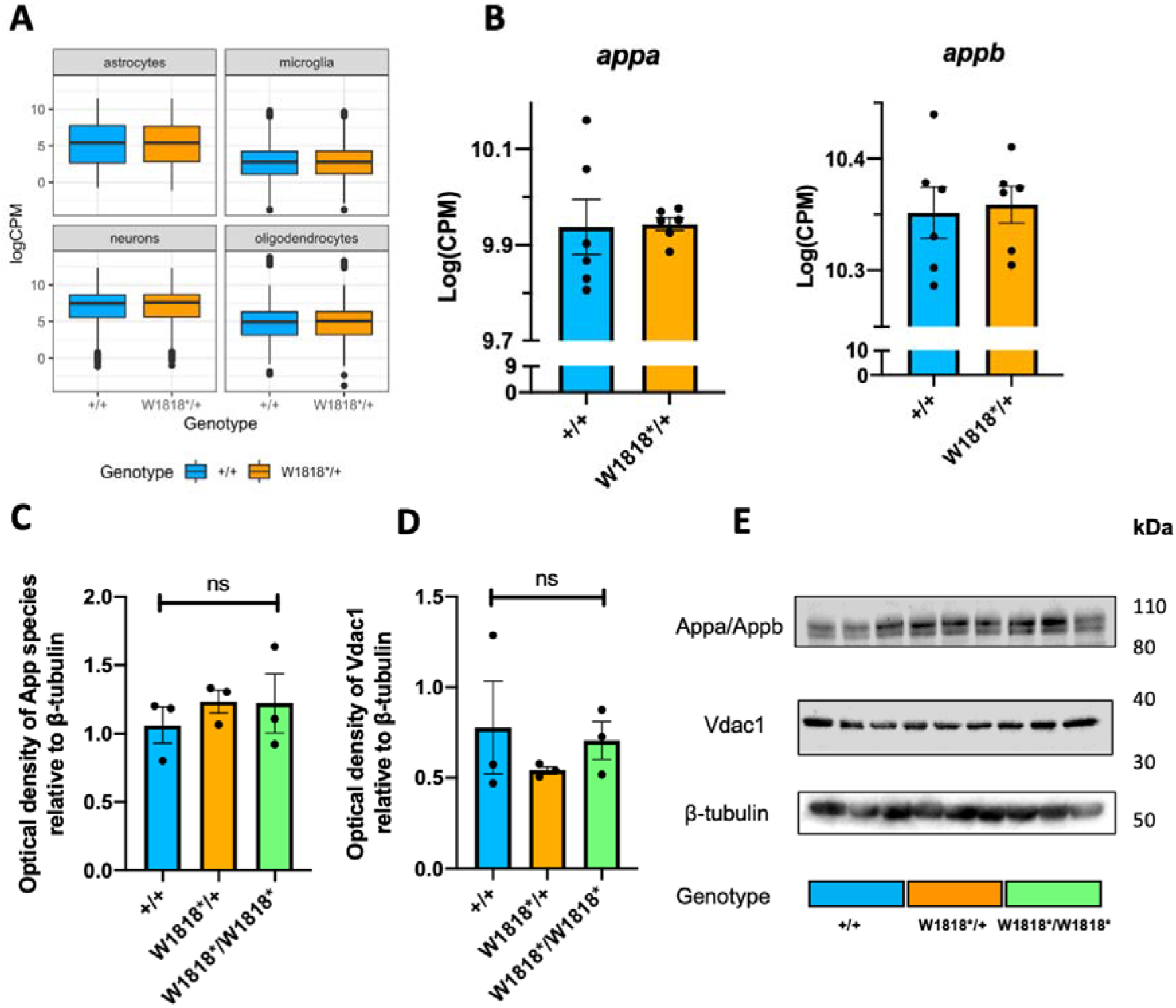
Changes to the transcriptome in W1818*/+ mutant brains are likely not due to changes in cell type proportions, altered expression of *appa/b*, or mitochondrial content. **A)** Distribution of expression (logCPM) of marker genes of astrocytes, microglia, neurons and oligodendrocytes in wild type (+/+) and heterozygous mutant (W1818*/+) brains. **B)** Expression (logCPM) of *APP* orthologues *appa* and *appb* in wild type and heterozygous mutant brains. **C)** Quantification of western blot analysis of expression of App proteins (Appa and Appb) in 6 month old wild type, heterozygous and homozygous mutant zebrafish brains. **D)** Quantification of western blot analysis of expression of the mitochondrial marker Vdac1 in 6 month old wild type, heterozygous and homozygous mutant zebrafish brains. **E)** Representative images of western immunoblots. Significance levels in **C)** and **D)** were determined by one-way ANOVA (ns, not significant).

Differences in transcription factor activity could also be driving the changes to gene expression observed in W1818*/+ brains. To explore this, we performed promotor motif enrichment analysis using *homer* (25) on the 100 most statistically significantly differentially expressed genes due to *sorl1* genotype. We identified that the promotor motif for hepatocyte nuclear factor 4 ⍰ (Hnf4⍰) is significantly over-represented in the promotors of the top 100 most DE genes (Bonferroni adjusted p-value = 0.04). Expression of *hnf4⍰* itself was low in these zebrafish brains and insufficient to be regarded as detectable in this RNA-seq experiment (< 0.75 cpm in at least 6 of the 12 RNA-seq libraries). However, examination of the logCPM values for *hnf4 ⍰* before filtering found that one mutant brain displayed far higher expression of *hnf4⍰* than the others and is clearly an outlier. A PCA on the expression of all genes which are predicted to contain a Hnf4*⍰*-binding motif (*HNF4ALPHA_Q6* gene set from the C3 category of *MSigDB*) revealed no obvious separation between W1818*/+ and +/+ samples, suggesting that *sorl1* genotype does not result in distinct expression patterns for these genes (**Additional File 6**).

In summary, heterozygosity for the W1818* mutation appears to have subtle effects on the expression of genes involved in energy production and protein translation in young adult zebrafish brains.

### Heterozygosity or homozygosity for W1818* does not appear to affect the abundance of App species, or mitochondrial mass

The most characterised cellular role of SORL1 protein is the sorting of APP throughout the endolysosomal system (reviewed in (11)). Therefore, we sought to investigate whether the W1818* mutation of *sorl1* causes changes in expression of the zebrafish forms of *APP*. In zebrafish, two paralogous genes show orthology with human *APP*: *appa* and *appb*.

Inspection of our RNA-seq data and did not reveal any differences in expression levels of *appa* or *appb* between heterozygous mutant and wild type brains (**Figure 4B**). We also used an antibody against human APP that cross-reacts with both zebrafish Appa and Appb proteins for immunoblot examination of App species in the brains of wild type, heterozygous and homozygous mutant zebrafish at 6 months of age (**Figure 4C**). We did not observe any significant differences between genotypes, supporting that the changes to gene expression are likely independent of changes to expression of *appa* or *appb*.

Finally, we hypothesised that in W1818*/+ brains, changes to expression of genes involved in oxidative phosphorylation could cause, or be due to, changes in cellular mitochondrial content. To explore this, we performed western immunoblotting against Vdac1, a protein highly abundant in the outer mitochondrial membrane, in lysates from entire zebrafish brains. We did not observe any significant effect of *sorl1* genotype on the levels of Vdac1, supporting that cellular mitochondrial content is unchanged between genotypes (**Figure 4D**).

## Discussion

We cannot access tissue from the living brains of *SORL1* mutation carriers for detailed molecular analysis. This restricts our ability to elucidate the early changes which eventually lead to AD and requires that we examine animal models instead. Our particular approach involves generating genetic models of EOfAD mutations in zebrafish which closely mimic the genetic state of human EOfAD (i.e. here we analysed the effects of a single, heterozygous mutation in the endogenous *sorl1* gene of zebrafish). This approach avoids potentially confounding assumptions such as that homozygous animals simply show more extreme phenotypes than heterozygotes, or that transgenic animals overexpressing mutant genes cause similar effects to endogenous, human mutations (assumptions commonly made in AD genetics research). We also performed our analyses when zebrafish are 6 months old and recently sexually mature. We regard this as the equivalent of early adulthood in humans. At this age changes to gene expression should reflect pathological processes (or the responses to these) occurring long before any cognitive deficits are thought to begin. Finally, analysis of large families of sibling fish raised in an identical environment (the same tank) reduces both genetic and environmental variation between samples (i.e. noise) and increases the likelihood of detecting subtle changes in gene expression due to heterozygosity for the EOfAD-like mutations.

In this study, we built on our previous analysis (14) investigating the effects of protein-truncating mutations in *sorl1* implicated in EOfAD. We generated a zebrafish model of the W1821* mutation in human *SORL1* (zebrafish *sorl1* W1818*). We showed that transcripts of the W1818* allele are likely subject to NMD (consistent with observations of coding sequence-truncating mutations in human *SORL1* (4)), and that W1818* transcripts cannot generate full-length Sorl1 protein. We did not observe any significant differences between the expression of *sorl1* transcripts in zebrafish brains under normoxia and hypoxia *in vivo*. This contrasts with the observations of Nishii et al. (15), using a hematopoietic stem cell line. These researchers saw increased SORL1 protein levels under hypoxia. However, a recent meta-analysis of RNA-seq datasets investigating transcriptional responses to hypoxia in both humans (128 datasets) and mice (52 datasets) showed that *SORL1* was seldom found to be differentially expressed under hypoxia. When *SORL1* was identified as DE, both up- and down-regulation was observed (26). Therefore, whether *SORL1* is regulated by hypoxia, and in which cell types this may occur, requires further investigation.

### Heterozygosity for EOfAD-like mutations in *sorl1* causes subtle changes in young adult brains

Transcriptome analysis of young adult, W1818*/+ mutant zebrafish brains relative to their +/+ siblings identified statistical evidence for two DE genes: *znf1148* and *prr15la*. Little or no information on the function of these genes can be found in the scientific literature. However, the candidate orthologue of *prr15la* in humans, *PRR15L*, was seen as associated with human intelligence in a meta-analysis of genome-wide association studies totalling 78,308 individuals (27). This low number of genes found to be DE is consistent with our previous analysis of another EOfAD-like mutation in *sorl1* (zebrafish *sorl1*^*V1482Afs*^ modelling human *SORL1*^*C1478\**^), for which heterozygosity gave rise to only one significantly DE (downregulated) gene: cytochrome c oxidase subunit 7A1 (*cox7a1*) (14). This gene appeared as slightly, but non-significantly, upregulated in W1818*/+ brains (logFC = 0.5, p = 0.2, p_FDR_ = 1) (**Additional File 4**). In contrast, analysis of an EOfAD-like mutation in the zebrafish orthologue of *PSEN1*, the gene most commonly mutated in EOfAD (28), detected 251 DE genes in the brains of young adult heterozygous zebrafish (29). Thus it appears that EOfAD mutations in *SORL1* cause less severe effects on cellular state compared to EOfAD mutations in *PSEN1*. This is more consistent with *SORL1*’s action as a LOAD genetic risk locus than as an EOfAD-causative locus. Campion and colleagues (30), noted that the ages of onset of AD in patients with mutations in *SORL1* are proximal to the arbitrary LOAD age threshold of 65 years of age (between 56 and 80 years, compared with *PSEN1* mutation carriers, which have an overall median age of onset of approximately 42 years (31)). Also, some *SORL1* mutation carriers with early onset AD carry additional LOAD genetic risk variants in *APOE, TREM2* and *ABCA7*, and/or have aged (> 66 years), unaffected siblings who carry the same *SORL1* mutation. Together, these observations, along with our *in vivo* transcriptomic studies, support that *SORL1* is more likely a genetic risk locus for LOAD than an EOfAD locus.

A promotor enrichment analysis on the top 100 genes which show the strongest signal for differential expression due to *sorl1* genotype identified the DNA-binding motif for hepatocyte nuclear factor 4 ⍰ (Hnf4⍰) as significantly enriched. *HNF4⍰*, a gene associated with maturity-onset diabetes of the young (MODY, a form of non-insulin-dependent diabetes mellitus (32)), is primarily expressed in the liver (reviewed in (33)) and has been reported to play roles in gluconeogenesis (34) and cholesterol homeostasis (35). Yamanishi and colleagues (36) reported that *HNF4⍰* is also expressed in the brains of mice where it appears to drive transcriptomic changes when the mice are housed under stressful conditions. Expression of *hnf4⍰* itself in the brains of our zebrafish was very low so that its transcripts were excluded during the pre-processing preceding our differentially gene expression analysis. However, inspection of the logCPM values from before exclusion of genes with low expression identified one W1818*/+ male mutant sample with a relatively higher expression of *hnf4⍰* (likely an outlier). Nevertheless, a PCA on the expression of all detectable genes predicted to contain the Hnf4⍰ DNA-binding motif (250 genes) did not reveal any convincing evidence of global dysregulation of Hnf4⍰-regulated genes in W1818*/+ brains.

### Changes to mitochondrial and ribosomal function are observed in young-adult, *sorl1* mutant zebrafish, and in human AD

Enrichment testing of all the detectable genes in the RNA-seq experiment identified subtle changes to expression of genes involved in energy production and protein translation. Reassuringly, we identified these processes as altered due to heterozygosity for another protein-truncating, EOfAD-like mutation in *sorl1*, V1482Afs, and for a putatively null mutation in *sorl1* in young adult zebrafish brains (14), suggesting that changes to mitochondrial and ribosomal function are an effect-in-common of EOfAD mutations in *SORL1*, and arise through a haploinsufficiency mechanism (i.e. due to decreased *sorl1* function).

Whether these mitochondrial and ribosomal functions are directly reliant on Sorl1 protein expression, or change as part of a homeostatic response to a deficiency of another, unknown, Sorl1-dependent function, is unclear. To our knowledge, the role of *SORL1* in the context of mitochondrial and ribosomal function has not been investigated in the scientific literature. However, it is well accepted that these processes are affected early in AD. Studies exploiting positron emission tomography with 2-[^18^F] fluoro-2-deoxy-d-glucose (FDG-PET) studies, (which measure glucose metabolism in the brains of living subjects), have shown that glucose metabolism gradually declines during the conversion from mild cognitive impairment (MCI) to AD, particularly in the parieto-temporal and posterior cingulate cortices (37-41). In our zebrafish model, genes encoding the components of the mitochondrial electron transport chain are mostly downregulated. Interestingly, the overall direction of change for genes involved in glycolysis and gluconeogenesis is up, supporting that a metabolic shift occurs from oxidative phosphorylation towards glycolysis as the primary source of energy for the brain. Additionally, changes to Hnf4⍰ activity may play a role in driving these changes to gluconeogenesis. However, further research is needed to test this possibility.

In W1818*/+ brains we observed downregulation of genes encoding ribosomal proteins. Ribosomes isolated from post-mortem MCI and AD patients show less capacity for protein synthesis (42-44) and increased levels of oxidised ribosomal RNA (rRNA) and bound ferrous iron (Fe^2+^) relative to control ribosomes (45). Oxidative stress within cells (such as due to increased redox active metals like Fe^2+^, or free radicals generated during normal mitochondrial respiration) oxidises rRNA and decreases ribosomal activity (46). Further investigation is needed to determine whether protein synthesis is actually affected in W1818*/+ brains.

We recently developed an enrichment analysis-based method to detect signs of iron dyshomeostasis in RNA-seq data. The method exploits gene sets encompassing genes that contain iron-responsive elements (IREs) in the 5’ or 3’ UTRs of their mRNAs (18). We did not observe any evidence for changes to expression of these gene sets, suggesting that intracellular iron (Fe^2+^) levels are not significantly changed in W1818*/+ brains relative to their +/+ siblings. This is consistent with our previous analysis, where we showed that heterozygosity for mutations in *sorl1* did not cause significant changes to expression of genes with IREs (but complete loss of wild type *sorl1* resulted in significant changes to expression of genes which IREs in the 3’ UTRs of their mRNAs) (14). Nevertheless, we cannot exclude that iron homeostasis is affected in the heterozygous mutant brains. Due to the nature of bulk RNA-seq, we may have failed to observe opposite directions of change in gene expression in different cell types.

In our previous analysis, we analysed a total of 24 fish which had one of four *sorl1* genotypes: +/+, EOfAD-like/+, null/+, or EOfAD-like/null (i.e. transheterozygous). In that experiment, 16 KEGG and HALLMARK gene sets were found to be altered significantly by heterozygosity for the EOfAD-like mutation (14). In contrast, the “two-genotype” analysis of this study found significant changes in only 6 gene sets. This suggests that the W1818* mutation (which models the human W1821* mutation in *SORL1*) appears to affect fewer cellular processes than the other EOfAD-like mutation (which models the C1478* mutation in human *SORL1*). Whether this reflects differing biological effects of the two mutations, or the greater statistical power of the “four-genotype” analysis (24 fish sequenced in the four-genotype analysis compared to 12 fish in the two-genotype analysis) is currently unclear. Future work may entail a four-genotype analysis of the W1818* mutation together with the null mutation in *sorl1* (R122Pfs) to compare directly whether W1818* resembles the loss-of-function, null mutation.

### No evidence that mutation of *sorl1* significantly changes expression of genes affecting the endo-lysosomal system in young adult zebrafish brains

We did not observe any gene sets involved in endolysosomal system function to be significantly altered in W1818*/+ brains, despite that SORL1 protein is thought to act, primarily, within the endolysosomal system (reviewed in (11)). Knupp and colleagues (47) showed that complete loss of *SORL1* in neurons (and not microglia) derived from human induced pluripotent stem cells (hiPSCs) resulted in early endosome enlargement, a phenomenon previously observed in post mortem EOfAD and LOAD brains (48-50). Our previous four-genotype analysis also did not reveal any direct evidence for dysregulation of genes in the endolysosomal system (14) although iron homeostasis appeared disturbed in brains lacking any wild type *sorl1* expression, and that might be due to a disturbance of cellular iron importation in which the endolysosomal system plays an important role (e.g. (51)). The results of our two independent zebrafish studies do not support that heterozygosity for mutations in *sorl1* affects the endolysosomal system at the level of gene regulation. Nevertheless, changes may be occurring in our mutant fish at the protein level without affecting the transcriptome. Also, as mentioned previously, humans heterozygous for *SORL1* mutations generally show ages of AD onset (if affected) close to the arbitrary late onset threshold of 65 years of age (4, 30). Therefore, it is possible that any endolysosomal defects in heterozygous W1818* zebrafish brains may not be observable until later ages.

In conclusion, we have provided evidence that mutation of *sorl1* affects mitochondrial and ribosomal function. Our bioinformatic analysis provides the basis for future experiments to elucidate the nature of these effects.

## Materials and Methods

### Zebrafish husbandry and animal ethics

All zebrafish used in this study were maintained in a recirculating water system on a 14 hour light/10 hour dark cycle, fed dry food in the morning and live brine shrimp in the afternoon. All zebrafish work was conducted under the auspices of the Animal Ethics Committee (permit numbers S-2017-089 and S-2017-073) and the Institutional Biosafety Committee of the University of Adelaide.

### Genome editing

To introduce mutations at the W1818 site in zebrafish *sorl1*, we used a TALEN pair designed and purchased from Zgenebio Biotech Inc. (Taipei City, Taiwan). The target genomic DNA sites (5’ to 3’) were ATGAGGTGGCGGTGTG (left) and GTAGGACTGTCTCCAT (right) (**Additional File 1**). The DNAs encoding the TALEN pairs were supplied in the pZGB2 vector, which were linearised with *Not*I (NEB, Ipswich, USA) before transcription of mRNA *in vitro* using the mMESSAGE mMACHINE T7 *in vitro* transcription kit following the manufacturer’s protocol (Invitrogen, Carlsbad, USA). The mRNAs encoding the TALEN pairs were diluted to a final concentration of 200 ng/µL each, and approximately 2-5 nL of the TALEN pair mRNA solution was injected into zebrafish embryos at the one cell stage. We eventually isolated a family of fish carrying an 11 nucleotide deletion in *sorl1*, W1818*, following the strategy described in (14). Briefly, injected embryos were raised until approximately three months of age, then were outcrossed to Tübingen strain fish of approximately the same age to generate F1 families. Since the injected fish are mosaic for any mutations at the W1818 site, we performed a T7 endonuclease I (T7EI) assay (NEB) on approximately 10 F1 embryos from a clutch to confirm whether any mutations had occurred in the germline of the injected parent. The PCR primers used to amplify the W1818 region for the T7EI assay have sequences (5’ – 3’) (F: TTAGGACCTCCTGTCAGCATTTCT and R: ACAAAATAAAAGTGTATGTGC). If the T7EI assay indicated that mutations were indeed in the germline, the remaining F1 embryos of the clutch were raised to adulthood. Sanger sequencing of the W1818 site was performed on genomic DNA extracted from fin biopsies of F1 adult fish to characterise any mutations (sequencing was performed by the Australian Genome Research Facility (AGRF, Adelaide, AUS)). Heterozygous mutant fish (W1818*/+) from the F1 family were pair mated with wild type (+/+) fish to generate F2 families. Heterozygous mutants from F2 families were in-crossed to produce F3 families containing homozygous mutant individuals.

### Hypoxia treatment and allele specific digital quantitative PCRs

Adult zebrafish were subjected to a hypoxic environment by placement in oxygen-depleted water for 3 hours (oxygen concentration of 6 ± 0.5 mg/L in normoxia and 1 ± 0.5mg/L in hypoxia). After treatment, fish were immediately sacrificed in a loose ice slurry and their brains removed. Total RNA was then extracted from whole zebrafish brains using the QIAGEN RNeasy⍰ Mini Kit (Qiagen, Venlo, Netherlands) according to the manufacturer’s protocol. Recovered RNA concentrations were estimated using a Nanodrop 2000 spectrophotometer (Thermo Fisher Scientific Inc, Waltham, Massachusetts, USA). cDNA was synthesized using random hexamers in the Superscript III First Strand Synthesis System (Invitrogen) according to the manufacturer’s instructions. 50 ng of each resulting cDNA sample was then used in allele-specific digital quantitative PCR as described in (52). To detect the two transcripts of *sorl1*, we used a W1818* mutation-specific forward primer with sequence 5’ GTGGCGGTGTGATGATGG 3’, and a wild type-specific primer with sequence 5’ TGGCGGTGTGGGCTCAC 3’. A common reverse primer sequence spanning the junction between exon 41 and 42 of *sorl1* (to reduce amplification from any genomic DNA carried over during RNA extraction) was 5’ GTAGAACACAGCGTACATCTCTGC 3’. Comparisons of mean transcript abundances per 50ng of brain-derived cDNA were made using a two-way ANOVA with Tukey’s post hoc test for multiple comparisons.

### Western blot analysis

Whole zebrafish brains were placed in 100 µL of 1X Complete Protease Inhibitor Cocktail (Roche Holding AG, Basel, Switzerland) in RIPA buffer (Sigma-Aldrich Corp. St. Louis, Missouri, USA), immediately homogenised using a handheld motorised pestle for 1 minute on ice, then incubated at 4°C for 2 hours with gentle rocking. Cellular debris was sedimented by centrifugation with a relative centrifugal force of 16,100 g for 10 minutes. 25 µL of 4X LDS buffer (Thermo Fisher Scientific) was added to the supernatant containing total protein, then samples were heated at 80°C for 20 minutes. Total protein concentrations were determined using the EZQ^®^ Protein Quantitation Kit (Molecular Probes, Inc. Eugene, OR, USA) following the manufacturers protocol. Samples were prepared for SDS-PAGE by adding 2.5 µL of 10X NuPAGE™ Sample Reducing Agent (Invitrogen) to 75 µg of total protein. The volume was increased to 25µL with 1X LDS buffer in RIPA buffer. Samples were heated at 80°C for 10 minutes and centrifuged briefly to sediment insoluble material. Supernatants containing the soluble proteins were subjected to SDS-PAGE on NuPAGE™ 4–12% Bis-Tris Protein Gels (Invitrogen) in the Mini Gel Tank and Blot Module Set (Thermo Fisher Scientific), using 1X NuPAGE™ MOPS SDS Running Buffer (Invitrogen). Resolved proteins were then transferred to PVDF membrane in 1X tris glycine SDS, 20% methanol. The PVDF membranes were blocked in 5% Western Blocking Reagent (Roche) then probed with primary and secondary antibodies. Antibody incubations were performed at either 1 hour at room temperature, or overnight at 4°C with gentle rocking. Antibody dilutions can be found in **Table 1**. Horse-radish peroxidase (HRP) signals were developed using Pierce™ ECL Western Blotting Substrate (Thermo Fisher Scientific) and imaged with a ChemiDoc™ MP Imaging System (Bio-Rad Laboratories, Hercules, California, USA). The intensities of the signals were measured using the Volume tool in Image Lab 5.1 (Bio-Rad Laboratories). The relative intensities of the signals were normalised to β-tubulin and compared across genotypes by one-way ANOVA tests. Original blot images can be found in **Additional File 8**.

**Table 1:** Antibodies used in this study.

| **Table 1: Antibodies used in this study.** |  |  |  |
| --- | --- | --- | --- |
| **Primary antibody** | **Dilution** | **Secondary Antibody** | **Dilution** |
| Goat Anti-SORL1 / LR11 (C Terminus)<br>(Aviva Systems Biology, San Diego, CA, USA) | 1:500 | Anti-goat IgG, HRP-conjugated Antibody<br>(Rockland Immunochemicals Inc. Limerick PA, USA) | 1:10,000 |
| Anti-VDAC<br>(Invitrogen) | 1:500 | Anti-rabbit IgG, HRP-conjugated Antibody<br>(Cell Signalling Technology, Danvers, MA, USA) | 1:10,000 |
| Anti-APP (clone 22C11)<br>(Sigma) | 1:3000 | Anti-mouse IgG, HRP-conjugated Antibody<br>(Rockland Immunochemicals) | 1:10,000 |
| Anti-β-Tubulin<br>(E7, deposited to the DSHB by Klymkowsky, M. (DSHB Hybridoma Product E7)) | 1:200 | Anti-mouse IgG, HRP-conjugated Antibody<br>(Rockland Immunochemicals) | 1:10,000 |

### RNA-seq analysis

We used a total of 12 fish in the RNA-seq analysis (i.e. n = 6 per genotype, with mostly equal males and females), based on a previous power calculation indicating that n = 6 would provide approximately 70% power to detect the majority of expressed transcripts in a zebrafish brain transcriptome at a fold-change > 2 and at a false discovery rate of 0.05 (data not shown). Whole heads were removed and preserved in approximately 600 µL of RNAlater (Thermo Fisher Scientific). Total RNA was extracted from the brains using the *mir*Vana miRNA isolation kit (Thermo Fisher Scientific) following the manufacturer’s protocol. Then, any genomic DNA carried over during the total RNA extraction was removed using the DNA-free™ Kit (Ambion, Waltham, USA). RNA was stabilised using RNAstable^®^ (Biomatrica, San Diego, CA, USA) following the manufacturers protocol to minimise RNA degradation. Briefly, RNA was resuspended in DEPC-treated water, applied directly to a RNAstable tube and dried with a vacuum concentrator. Dried RNA was sent to Novogene (Hong King, China) for cDNA library synthesis and RNA-sequencing. The cDNA libraries were generated using NEBNext Ultra RNA Library Prep Kit for Illumina (NEB) and RNA was sequenced using the Illumina Novaseq PE150 platform.

Demultiplexed paired-end fastq files (with adapter sequences removed) were supplied by Novogene. We first inspected the quality of the raw data by *fastQC* and *ngsReports* (53). We then performed a pseudo alignment using *kallisto* (54) version 0.43.1, in paired end mode specifying 50 bootstraps. The index file for *kallisto* was generated according to the zebrafish transcriptome (primary assembly of GRCz11, Ensembl release 96), with the sequences for unspliced transcripts additionally included to enable confirmation of genotypes. We imported the transcript counts estimated by *kallisto* for analysis using *R* (55), using the *catchKallisto* function of the package *edgeR* (56). To obtain gene level counts, we summed the counts of all transcripts arising from a single gene. Normalisation factors were calculated using the trimmed mean of M-values (TMM) method (57). Low expressed genes are statistically uninformative as they provide little evidence for differential expression, and we considered genes to be detectably expressed if they contained at least contained a logCPM of 0.75 in at least 6 of the 12 RNA-seq libraries. After selecting detectable genes, library sizes ranged from 13,050,237 to 50,560,749. Although these library sizes varied considerably, a correlation between library size and the first two principal components was not observed, supporting that variation due to library size does not contribute to the two largest sources of variation in this dataset (**Additional File 7**).

We performed an initial differential expression analysis using the exact test function of *edgeR* (56). A design matrix was specified with an intercept for each brain sample, and *sorl1* genotype (W1818*/+) as the common difference. Only one differentially expressed gene was identified in this analysis (data not shown). Therefore, to assist with identification of dysregulated genes due to *sorl1* genotype, we removed one factor of unwanted variation using the *RUVg* method of the package *RUVSeq* (58). For *RUVg*, we set *k* = 1, and the negative control genes as the least 10,000 differentially expressed genes (i.e. largest p-value) in the initial differential expression test. The W_1 offset term generated was then included in the design matrix for an additional differential expression test using a generalised linear model and likelihood ratio tests with *edgeR* (56, 59). We considered genes differentially expressed if their FDR adjusted p-value was less than 0.05.

Gene set enrichment testing was performed using three different algorithms: *fry* (19), *camera* (21) and *GSEA* (17, 20). Since each method gave different levels of significance, we combined the raw p-values from each method by calculating the harmonic mean p-value (22). We considered a gene set to be significantly altered if the FDR adjusted harmonic mean p-value was less than 0.05. The gene sets we tested in this study were the *KEGG* (60), *HALLMARK* (16) and the zebrafish iron-responsive element (18) gene sets. The *KEGG* and *HALLMARK* gene sets for zebrafish were obtained from *MSigDB* (16) using *msigdbR* (61). To perform promoter motif enrichment analysis, we used *homer* (25) as described in (62) on the top 100 most statistically significant differentially expressed genes due to *sorl1* genotype when including the W_1 covariate in the design matrix.

Data visualisation was performed using the packages *ggplot2* (63), *pheatmap* (64), and *upsetR* (65).

## Declarations

### Ethics approval and consent to participate

All zebrafish work was conducted under the auspices of the University of Adelaide Animal Ethics Committee (permit numbers: S-2017-073 and S-2017-089) and Institutional Biosafety Committee (permit number 15037).

## Consent for publication

Not applicable.

## Availability of data and materials

The raw fastq files and the output from *catchKallisto* (i.e. the raw transcript abundances) have been deposited in Gene Expression Omnibus database (GEO) under Accession Number GSE156167. All code to reproduce the RNA-seq analysis can be found at https://github.com/karissa-b/sorl1_w1818x_6m.

## Authors’ contributions

KB performed all experimental and bioinformatic analysis and drafted the manuscript. SP supervised and provided advice on the bioinformatic analysis and MN and ML supervised all zebrafish work. All authors read and contributed to the final manuscript.

## Acknowledgements

This work was supported with supercomputing resources provided by the Phoenix HPC service at the University of Adelaide.

The authors would like to acknowledge Dr Seyyed Hani Moussavi Nik for obtaining part of the funding for this work, and his assistance in the genome editing to generate the W1818* mutant line of zebrafish, Ms Nhi Hin for providing the zebrafish iron-responsive element (IRE) gene sets and her valuable advice and discussions on the bioinformatic analysis, and Dr Zijing Zhou for his assistance in performing the western blots.

## Additional Files

**Additional File 1:**
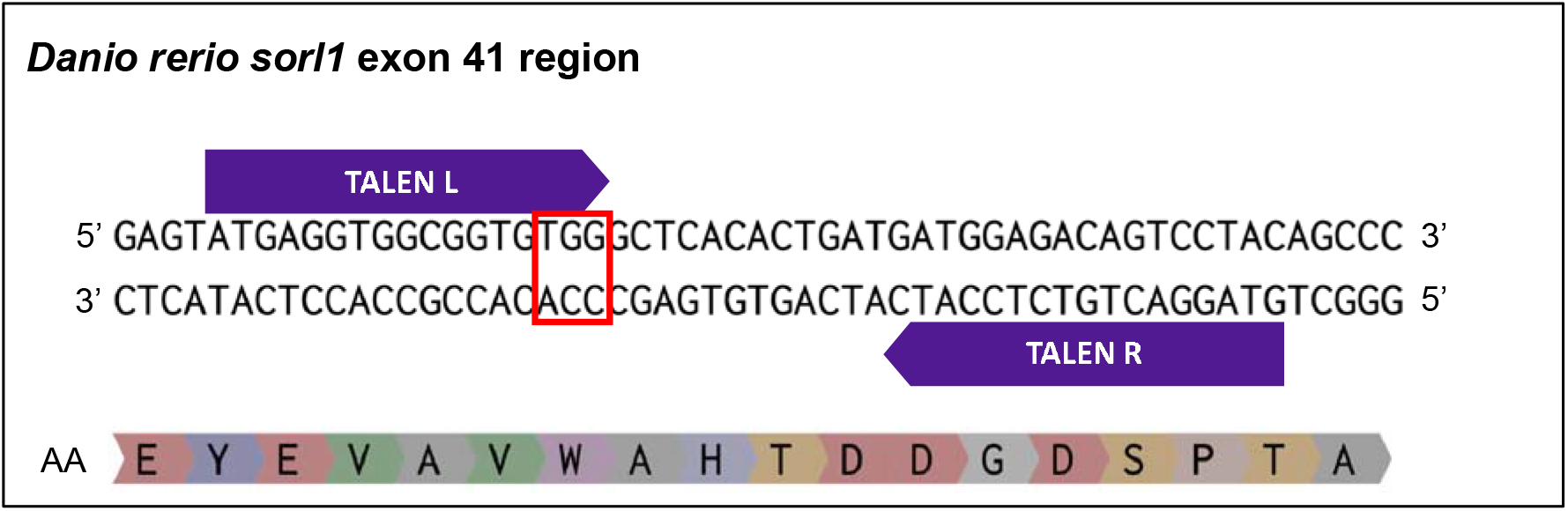
Genome editing of *sorl1* exon 41. A 55 nucleotide section of zebrafish *sorl1* exon 41 (ENSDARE00000237070). Both the sense, anti-sense and the translated (AA) sequences are indicated. The TALEN recognition sites are indicated by purple arrows, and the W1818 site is highlighted by the red box.

**Additional File 2:**
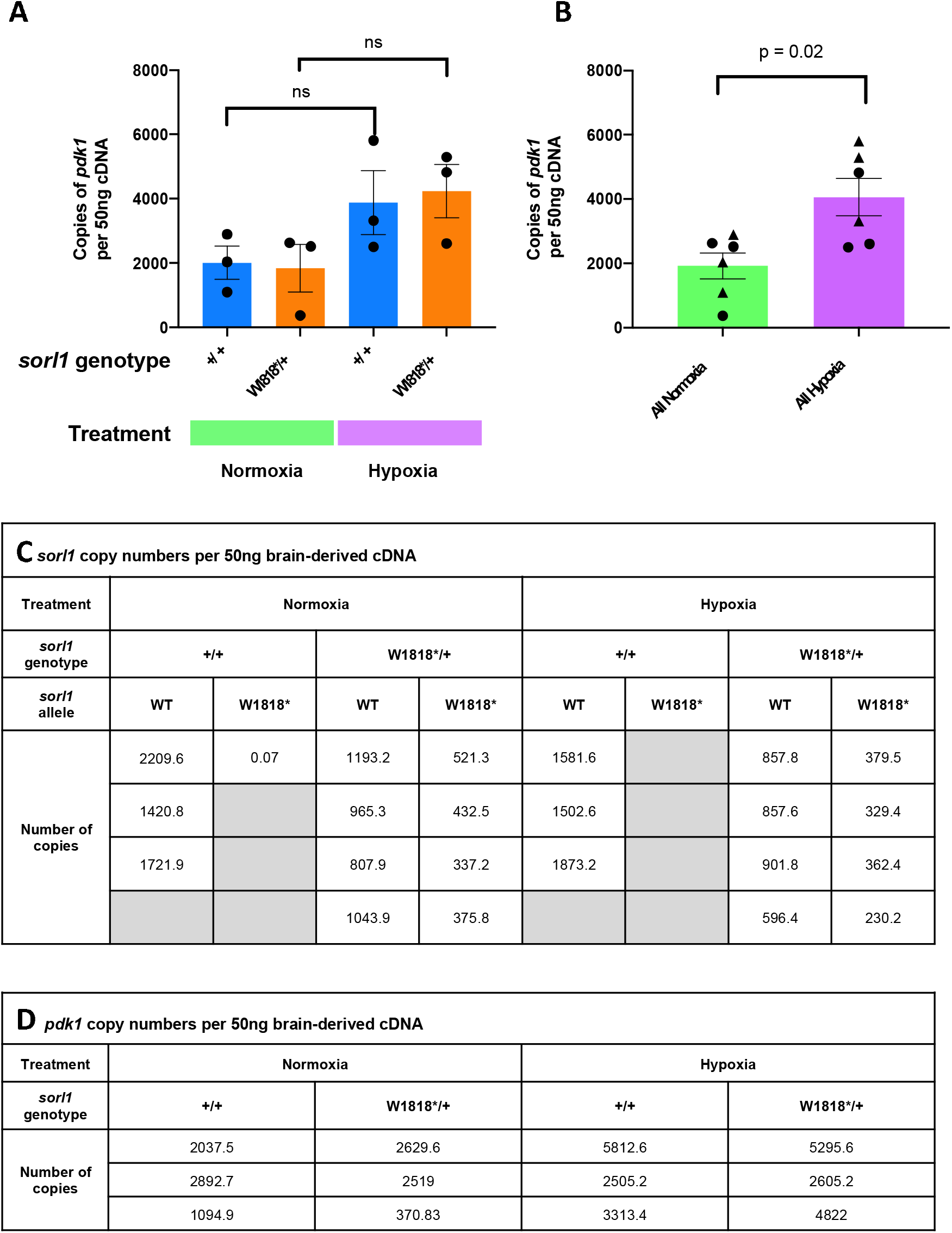
Expression of *sorl1* and *pdk1* in young adult zebrafish brains exposed to hypoxia. **A)** Mean ± SEM copies of pyruvate dehydrogenase kinase (*pdk1*) per 50ng of brain derived cDNA of zebrafish either heterozygous for the W1818* mutation in *sorl1* (W1818*/+), or their wild type siblings (+/+) and exposed to either normoxia or hypoxia. Expression of *pdk1* appeared to increase after exposure to acute hypoxia in each *sorl1* genotype. However, these comparisons did not reach statistical significance (one-way ANOVA with Dunnet’s post-hoc test). **B)** Re-analysis of the data in **A**, where data points were combined by treatment. A Student’s *t*-test identified a significant increase in *pdk1* expression. +/+ samples are indicated by triangles, and W1818*/+ samples are indicated by circles. **C)** Expression levels of *sorl1* transcripts per 50 ng of brain-derived cDNA under normoxia and hypoxia. Numbers represent the copy numbers on the QuantStudio™ 3D Digital PCR 20K chip after processing within QuantStudio™ 3D AnalysisSuite Cloud Software (version 3.0, Applied Biosystems, Thermo Fisher Scientific). Note that only one digital PCR was performed to detect the W1818* allele of *sorl1* in +/+ brains under normoxia to confirm that the W1818* mutation specific primers did not amplify from the wild type (WT) sequence of *sorl1*. **D)** Expression levels of *pdk1* transcripts per 50 ng of brain-derived cDNA under normoxia and hypoxia.

**Additional File 3:**
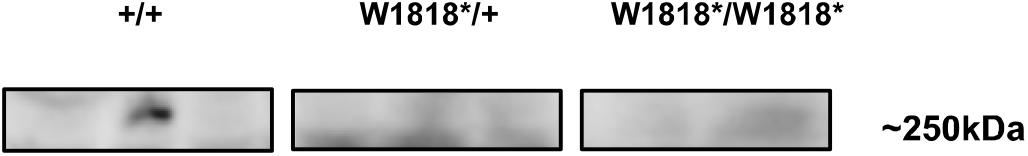
Expression of Sorl1 protein in young adult zebrafish brains. Western immunoblot against C-terminal sequences of Sorl1 protein in 6 month old wild type, heterozygous mutant and homozygous mutant sibling brains. Loss of the signal at approximately 250 kDa supports that the W1818* mutation results in disruption of translation of Sorl1 protein.

**Additional File 4: Full differential expression analysis results**

DGE_results_RUV.csv

The full results for the differential gene expression analysis using edgeR and including the W_1 covariate from *RUVseq* in the design matrix.

**Additional file 5:**
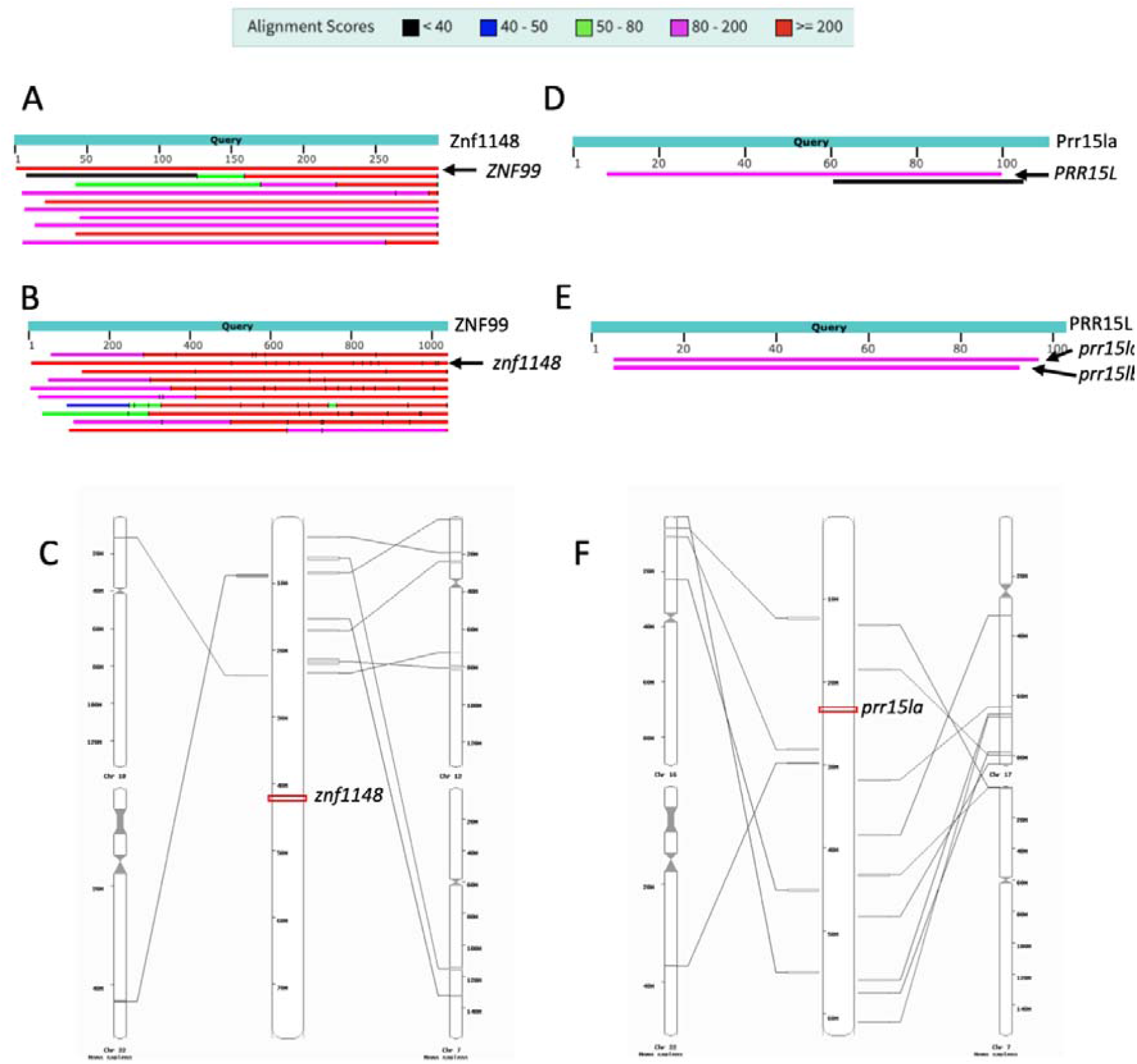
BLAST and analysis of synteny of the gene differentially expressed due to heterozygosity for the W1818* mutation of *sorl1*. **A)** Graphical summary of the top 10 hits of a tblastn search using the protein sequence of Znf1148 (ENSDART00000163798.2) against the human genome GRCh38.p13. The best hit corresponds to *ZNF99* on human chromosome 19 (chr19). **B)** Graphical summary of the best 10 hits of a tblastn search using the protein sequence of human ZNF99 (ENST00000397104.5) against the zebrafish genome GRCz11. As a reverse viewpoint to **A**, the second hit refers to *znf1148*. **C)** Analysis of synteny between zebrafish chromosome 4 (chr4) and the human genome. The location of *znf1148* is shown in red. No conservation of synteny is observed in the human genome with the *znf1148* region. **D)** Graphical summary of a tblastn search using the protein sequence of zebrafish Prr15la (ENSDART00000134723.2) against the human genome GRCh38.p13. The top hit corresponds to *PRR15L* on human chromosome 17 (chr17). **E)** Graphical summary of the top 10 hits of a tblastn search using the protein sequence of PRR15L (ENST00000300557.3) against the zebrafish genome GRCz11. The only significant homology is to the paralogues *prr15la* and *prr15lb*. **F)** Analysis of synteny between zebrafish chromosome 3 (chr3) and the human genome. The location of *prr15la* is shown in red. No conservation of synteny is observed in the human genome with the *prr15la* region. BLAST searches and analysis of synteny were performed using the Ensembl web site (https://m.ensembl.org/).

**Additional File 6:**
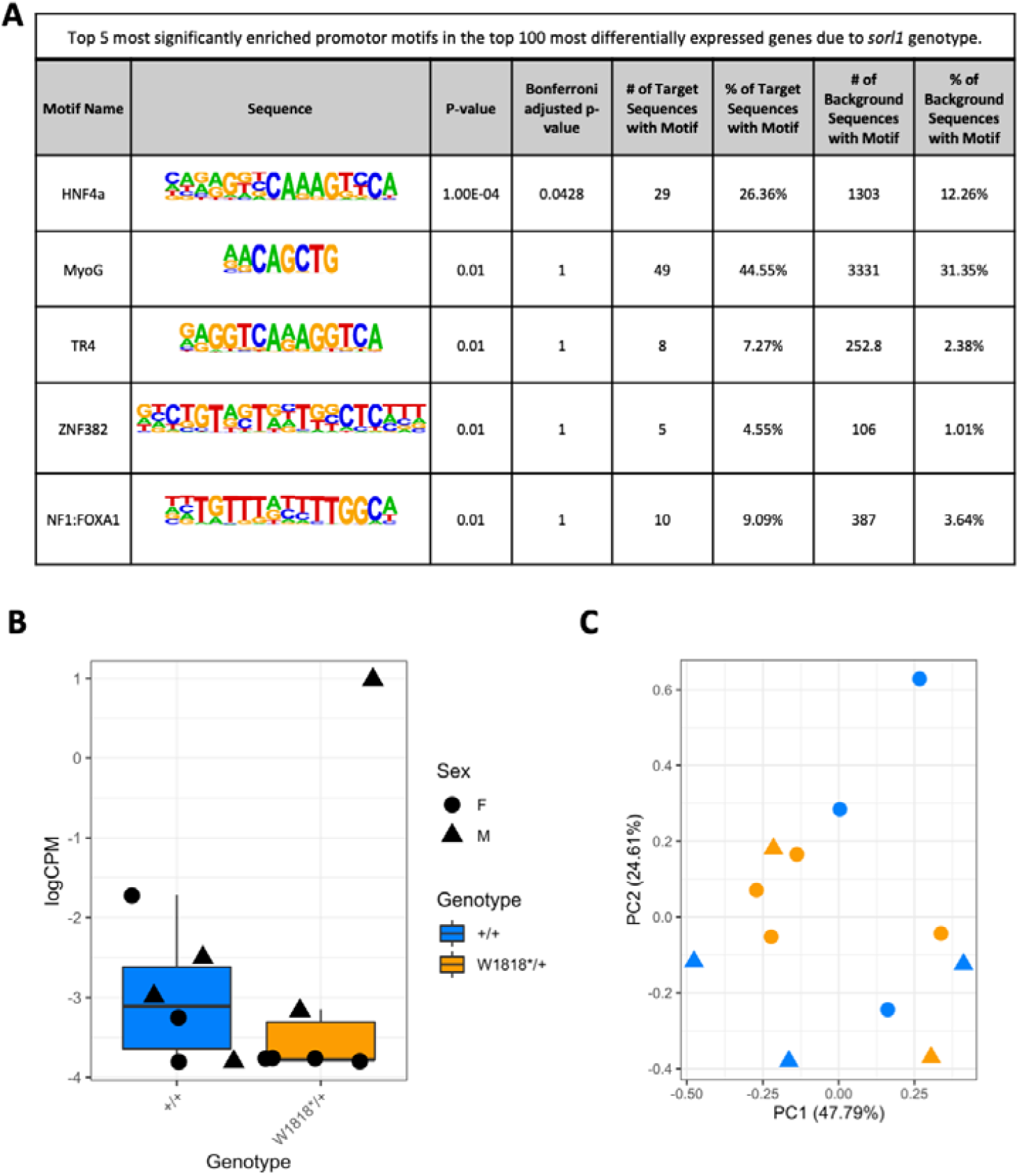
Promotor motif analysis using homer. **A)** The top 5 most significantly enriched motifs in the promoters of the top 100 most statistically significant gene differentially expressed due to *sorl1* genotype, relative to all genes detected in the RNA-seq experiment. **B)** Expression of *hnf4*⍰ before filtering lowly expressed genes in the RNA-seq dataset. **C)** Principal component (PC) analysis plot of the *RUVseq* adjusted expression values of genes containing a motif for binding of Hnf4⍰ (*HNF4ALPHA_Q6* gene set from *MSigDB*, C3 category, TFT:TFT_Legacy subcategory).

**Additional file 7:**
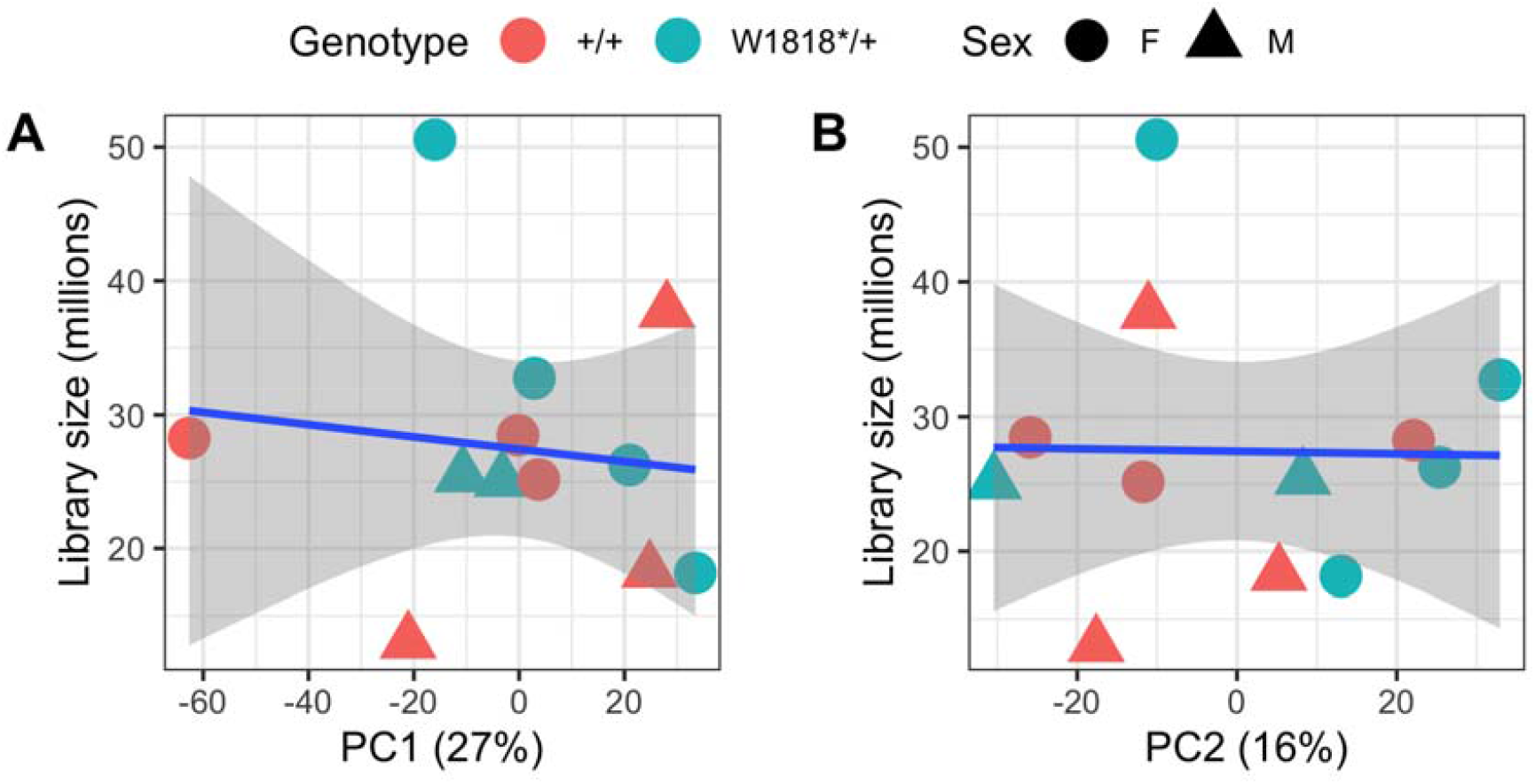
PC1 and PC2 do not correlate with library size. Plot of the **A)** principal component 1 (PC1) and **B)** PC2 value against the size of each RNA-seq library. The blue lines indicates a linear model regression lines with standard error shading. No correlation is observed, suggesting that the varying library sizes does not contribute overly to the two largest sources of variation in this dataset.

**Additional File 8:**
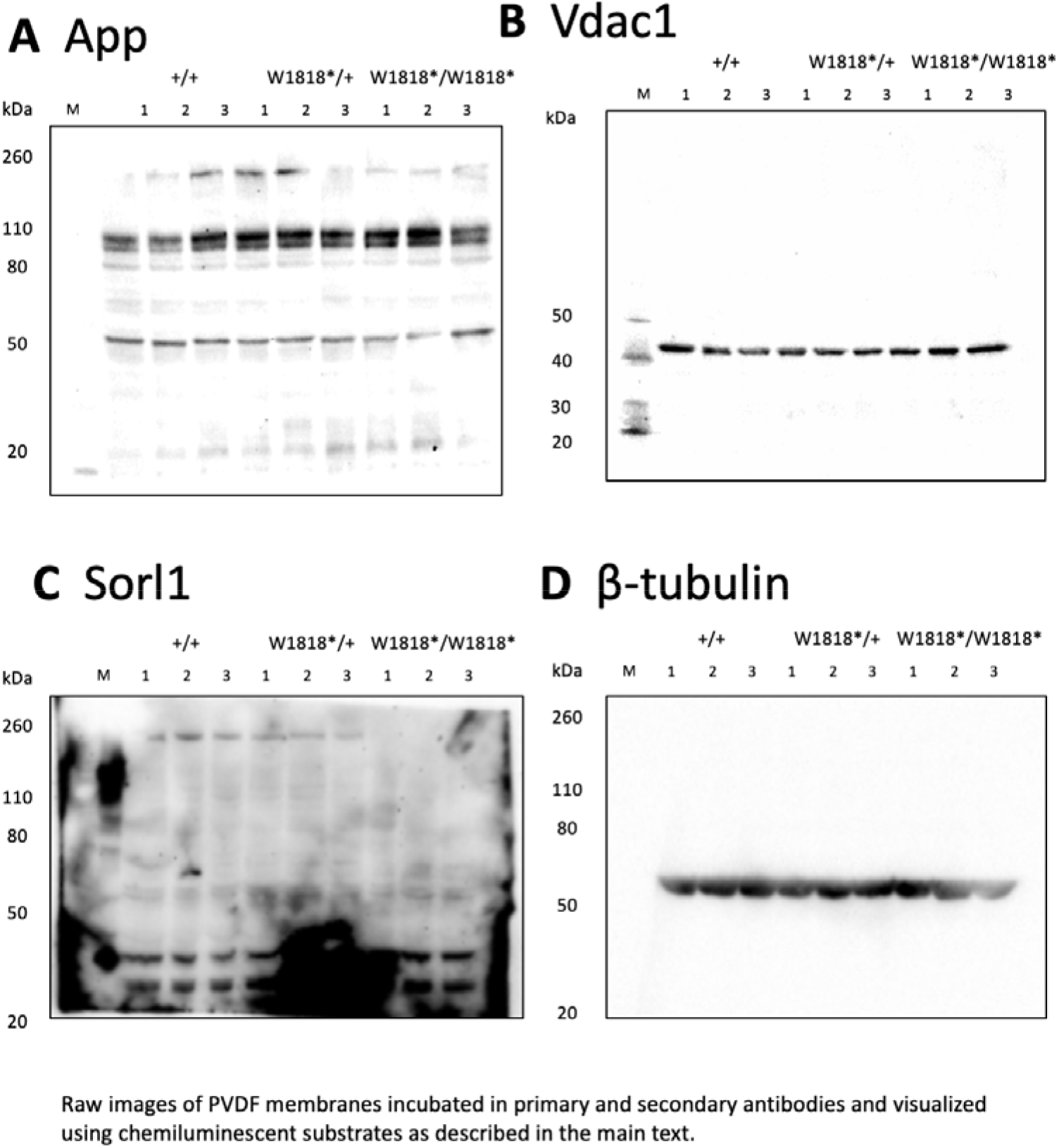
Original images of western blot membranes.

## References

1. Kunkle BW, Grenier-Boley B, Sims R, Bis JC, Damotte V, Naj AC, et al. Genetic meta-analysis of diagnosed Alzheimer’s disease identifies new risk loci and implicates Aβ, tau, immunity and lipid processing. Nature Genetics. 2019;51(3):414–30.

2. Lambert J-C, Ibrahim-Verbaas CA, Harold D, Naj AC, Sims R, Bellenguez C, et al. Meta-analysis of 74,046 individuals identifies 11 new susceptibility loci for Alzheimer’s disease. Nature Genetics. 2013;45:1452.

3. Masters CL, Bateman R, Blennow K, Rowe CC, Sperling RA, Cummings JL. Alzheimer’s disease. Nature Reviews Disease Primers. 2015;1(1):15056.

4. Pottier C, Hannequin D, Coutant S, Rovelet-Lecrux A, Wallon D, Rousseau S, et al. High frequency of potentially pathogenic SORL1 mutations in autosomal dominant early-onset Alzheimer disease. Molecular psychiatry. 2012;17(9):875–9.

5. Thonberg H, Chiang H-H, Lilius L, Forsell C, Lindström A-K, Johansson C, et al. Identification and description of three families with familial Alzheimer disease that segregate variants in the SORL1 gene. Acta Neuropathologica Communications. 2017;5(1):43.

6. Verheijen J, Van den Bossche T, van der Zee J, Engelborghs S, Sanchez-Valle R, Lladó A, et al. A comprehensive study of the genetic impact of rare variants in SORL1 in European early-onset Alzheimer’s disease. Acta Neuropathologica. 2016;132(2):213–24.

7. Lee JH, Cheng R, Schupf N, Manly J, Lantigua R, Stern Y, et al. The association between genetic variants in SORL1 and Alzheimer disease in an urban, multiethnic, community-based cohort. Archives of neurology. 2007;64(4):501–6.

8. Reitz C, Cheng R, Rogaeva E, Lee JH, Tokuhiro S, Zou F, et al. Meta-analysis of the association between variants in SORL1 and Alzheimer disease. Archives of neurology. 2011;68(1):99–106.

9. Rogaeva E, Meng Y, Lee JH, Gu Y, Kawarai T, Zou F, et al. The neuronal sortilin-related receptor SORL1 is genetically associated with Alzheimer disease. Nat Genet. 2007;39(2):168–77.

10. Wen Y, Miyashita A, Kitamura N, Tsukie T, Saito Y, Hatsuta H, et al. SORL1 is genetically associated with neuropathologically characterized late-onset Alzheimer’s disease. Journal of Alzheimer’s disease : JAD. 2013;35(2):387–94.

11. Barthelson K, Newman M, Lardelli M. Sorting Out the Role of the Sortilin-Related Receptor 1 in Alzheimer’s Disease. Journal of Alzheimer’s Disease Reports. 2020;Preprint:1–18.

12. Caglayan S, Takagi-Niidome S, Liao F, Carlo A-S, Schmidt V, Burgert T, et al. Lysosomal Sorting of Amyloid-β by the SORLA Receptor Is Impaired by a Familial Alzheimer’s Disease Mutation. Science Translational Medicine. 2014;6(223):223ra20–ra20.

13. Yajima R, Tokutake T, Koyama A, Kasuga K, Tezuka T, Nishizawa M, et al. ApoE-isoform-dependent cellular uptake of amyloid-beta is mediated by lipoprotein receptor LR11/SorLA. Biochemical and biophysical research communications. 2015;456(1):482–8.

14. Barthelson K, Pederson SM, Newman M, Lardelli M. Brain transcriptome analysis reveals subtle effects on mitochondrial function and iron homeostasis of mutations in the SORL1 gene implicated in early onset familial Alzheimer’s disease. bioRxiv. 2020:2020.07.17.207787.

15. Nishii K, Nakaseko C, Jiang M, Shimizu N, Takeuchi M, Schneider WJ, et al. The Soluble Form of LR11 Protein Is a Regulator of Hypoxia-induced, Urokinase-type Plasminogen Activator Receptor (uPAR)-mediated Adhesion of Immature Hematological Cells. Journal of Biological Chemistry. 2013;288(17):11877–86.

16. Liberzon A, Birger C, Thorvaldsdóttir H, Ghandi M, Mesirov Jill P, Tamayo P. The Molecular Signatures Database Hallmark Gene Set Collection. Cell Systems. 2015;1(6):417–25.

17. Subramanian A, Tamayo P, Mootha VK, Mukherjee S, Ebert BL, Gillette MA, et al. Gene set enrichment analysis: A knowledge-based approach for interpreting genome-wide expression profiles. Proceedings of the National Academy of Sciences. 2005;102(43):15545.

18. Hin N, Newman M, Pederson SM, Lardelli MM. Iron Responsive Element (IRE)-mediated responses to iron dyshomeostasis in Alzheimer’s disease. bioRxiv. 2020:2020.05.01.071498.

19. Wu D, Lim E, Vaillant F, Asselin-Labat M-L, Visvader JE, Smyth GK. ROAST: rotation gene set tests for complex microarray experiments. Bioinformatics. 2010;26(17):2176–82.

20. Sergushichev AA. An algorithm for fast preranked gene set enrichment analysis using cumulative statistic calculation. bioRxiv. 2016:060012.

21. Wu D, Smyth GK. Camera: a competitive gene set test accounting for inter-gene correlation. Nucleic acids research. 2012;40(17):e133–e.

22. Wilson DJ. The harmonic mean p-value for combining dependent tests.Proceedings of the National Academy of Sciences. 2019;116(4):1195.

23. Cahoy JD, Emery B, Kaushal A, Foo LC, Zamanian JL, Christopherson KS, et al. A transcriptome database for astrocytes, neurons, and oligodendrocytes: a new resource for understanding brain development and function. The Journal of neuroscience : the official journal of the Society for Neuroscience. 2008;28(1):264–78.

24. Oosterhof N, Holtman IR, Kuil LE, van der Linde HC, Boddeke EWGM, Eggen BJL, et al. Identification of a conserved and acute neurodegeneration-specific microglial transcriptome in the zebrafish. Glia. 2017;65(1):138–49.

25. Heinz S, Benner C, Spann N, Bertolino E, Lin YC, Laslo P, et al. Simple combinations of lineage-determining transcription factors prime cis-regulatory elements required for macrophage and B cell identities. Molecular cell. 2010;38(4):576–89.

26. Bono H, Hirota K. Meta-Analysis of Hypoxic Transcriptomes from Public Databases. Biomedicines. 2020;8(1).

27. Sniekers S, Stringer S, Watanabe K, Jansen PR, Coleman JRI, Krapohl E, et al. Genome-wide association meta-analysis of 78,308 individuals identifies new loci and genes influencing human intelligence. Nature Genetics. 2017;49(7):1107–12.

28. Cruts M, Theuns J, Van Broeckhoven C. Locus-specific mutation databases for neurodegenerative brain diseases. Human Mutation. 2012;33(9):1340–4.

29. Newman M, Hin N, Pederson S, Lardelli M. Brain transcriptome analysis of a familial Alzheimer’s disease-like mutation in the zebrafish presenilin 1 gene implies effects on energy production. Molecular Brain. 2019;12(1).

30. Campion D, Charbonnier C, Nicolas G. SORL1 genetic variants and Alzheimer disease risk: a literature review and meta-analysis of sequencing data. Acta Neuropathologica. 2019.

31. Ryman DC, Acosta-Baena N, Aisen PS, Bird T, Danek A, Fox NC, et al. Symptom onset in autosomal dominant Alzheimer disease: A systematic review and meta-analysis. Neurology. 2014;83(3):253–60.

32. Hani EH, Suaud L, Boutin P, Chèvre JC, Durand E, Philippi A, et al. A missense mutation in hepatocyte nuclear factor-4 alpha, resulting in a reduced transactivation activity, in human late-onset non-insulin-dependent diabetes mellitus. J Clin Invest. 1998;101(3):521–6.

33. Cereghini S. Liver-enriched transcription factors and hepatocyte differentiation. The FASEB Journal. 1996;10(2):267–82.

34. Rhee J, Inoue Y, Yoon JC, Puigserver P, Fan M, Gonzalez FJ, et al. Regulation of hepatic fasting response by PPARγ coactivator-1α (PGC-1): Requirement for hepatocyte nuclear factor 4α in gluconeogenesis. Proceedings of the National Academy of Sciences. 2003;100(7):4012.

35. Yin L, Ma H, Ge X, Edwards PA, Zhang Y. Hepatic hepatocyte nuclear factor 4α is essential for maintaining triglyceride and cholesterol homeostasis. Arterioscler Thromb Vasc Biol. 2011;31(2):328–36.

36. Yamanishi K, Doe N, Sumida M, Watanabe Y, Yoshida M, Yamamoto H, et al. Hepatocyte Nuclear Factor 4 Alpha Is a Key Factor Related to Depression and Physiological Homeostasis in the Mouse Brain. PLOS ONE. 2015;10(3):e0119021.

37. Mosconi L. Brain glucose metabolism in the early and specific diagnosis of Alzheimer’s disease. FDG-PET studies in MCI and AD. Eur J Nucl Med Mol Imaging. 2005;32(4):486–510.

38. Mosconi L, Mistur R, Switalski R, Tsui WH, Glodzik L, Li Y, et al. FDG-PET changes in brain glucose metabolism from normal cognition to pathologically verified Alzheimer’s disease. Eur J Nucl Med Mol Imaging. 2009;36(5):811–22.

39. Drzezga A, Lautenschlager N, Siebner H, Riemenschneider M, Willoch F, Minoshima S, et al. Cerebral metabolic changes accompanying conversion of mild cognitive impairment into Alzheimer’s disease: a PET follow-up study. Eur J Nucl Med Mol Imaging. 2003;30(8):1104–13.

40. Minoshima S, Giordani B, Berent S, Frey KA, Foster NL, Kuhl DE. Metabolic reduction in the posterior cingulate cortex in very early Alzheimer’s disease. Annals of Neurology. 1997;42(1):85–94.

41. Iturria-Medina Y, Sotero R, Toussaint P, Mateos-Pérez J, Evans A, Initiative AsDN. Early role of vascular dysregulation on late-onset Alzheimer’s disease based on multifactorial data-driven analysis. Nature Communications. 2016;7.

42. Ding Q, Markesbery WR, Chen Q, Li F, Keller JN. Ribosome dysfunction is an early event in Alzheimer’s disease. The Journal of neuroscience : the official journal of the Society for Neuroscience. 2005;25(40):9171–5.

43. Ding Q, Markesbery WR, Cecarini V, Keller JN. Decreased RNA, and Increased RNA Oxidation, in Ribosomes from Early Alzheimer’s Disease. Neurochemical Research. 2006;31(5):705–10.

44. Langstrom NS, Anderson JP, Lindroos HG, Winbland B, Wallace WC. Alzheimer’s disease-associated reduction of polysomal mRNA translation. Molecular Brain Research. 1989;5(4):259–69.

45. Honda K, Smith MA, Zhu X, Baus D, Merrick WC, Tartakoff AM, et al. Ribosomal RNA in Alzheimer Disease Is Oxidized by Bound Redox-active Iron. Journal of Biological Chemistry. 2005;280(22):20978–86.

46. Willi J, Küpfer P, Evéquoz D, Fernandez G, Katz A, Leumann C, et al. Oxidative stress damages rRNA inside the ribosome and differentially affects the catalytic center. Nucleic Acids Research. 2018;46(4):1945–57.

47. Knupp A, Mishra S, Martinez R, Braggin JE, Szabo M, Kinoshita C, et al. Depletion of the AD Risk Gene SORL1 Selectively Impairs Neuronal Endosomal Traffic Independent of Amyloidogenic APP Processing. Cell Reports. 2020;31(9):107719.

48. Cataldo AM, Petanceska S, Peterhoff CM, Terio NB, Epstein CJ, Villar A, et al. App gene dosage modulates endosomal abnormalities of Alzheimer’s disease in a segmental trisomy 16 mouse model of down syndrome. The journal of neuroscience. 2003;23(17):6788–92.

49. Cataldo AM, Peterhoff CM, Schmidt SD, Terio NB, Duff K, Beard M, et al. Presenilin mutations in familial Alzheimer disease and transgenic mouse models accelerate neuronal lysosomal pathology. Journal of Neuropathology & Experimental Neurology. 2004;63(8):821–30.

50. Cataldo AM, Peterhoff CM, Troncoso JC, Gomez-Isla T, Hyman BT, Nixon RA. Endocytic pathway abnormalities precede amyloid beta deposition in sporadic Alzheimer’s disease and Down syndrome: differential effects of APOE genotype and presenilin mutations. The American journal of pathology. 2000;157(1):277–86.

51. Yambire KF, Rostosky C, Watanabe T, Pacheu-Grau D, Torres-Odio S, Sanchez-Guerrero A, et al. Impaired lysosomal acidification triggers iron deficiency and inflammation in vivo. Elife. 2019;8.

52. Jiang H, Newman M, Lardelli M. The zebrafish orthologue of familial Alzheimer’s disease gene PRESENILIN 2 is required for normal adult melanotic skin pigmentation. PLOS ONE. 2018;13(10):e0206155.

53. Ward CM, To TH, Pederson SM. ngsReports: a Bioconductor package for managing FastQC reports and other NGS related log files. Bioinformatics. 2020;36(8):2587–8.

54. Bray NL, Pimentel H, Melsted P, Pachter L. Near-optimal probabilistic RNA-seq quantification. Nature Biotechnology. 2016;34:525.

55. Team RC. R: A language and environment for statistical computing. R Foundation for Statistical Computing, Vienna, Austria. 2019.

56. Robinson MD, McCarthy DJ, Smyth GK. edgeR: a Bioconductor package for differential expression analysis of digital gene expression data. Bioinformatics. 2009;26(1):139–40.

57. Robinson MD, Oshlack A. A scaling normalization method for differential expression analysis of RNA-seq data. Genome Biology. 2010;11(3):R25.

58. Risso D, Ngai J, Speed TP, Dudoit S. Normalization of RNA-seq data using factor analysis of control genes or samples. Nature Biotechnology. 2014;32(9):896–902.

59. McCarthy DJ, Chen Y, Smyth GK. Differential expression analysis of multifactor RNA-Seq experiments with respect to biological variation. Nucleic Acids Research. 2012;40(10):4288–97.

60. Kanehisa M, Goto S. KEGG: kyoto encyclopedia of genes and genomes. Nucleic acids research. 2000;28(1):27–30.

61. Dolgalev I. msigdbr: MSigDB Gene Sets for Multiple Organisms in a Tidy Data Format. 7.1.1 ed 2020. p. R package.

62. Hin N, Newman M, Kaslin J, Douek AM, Lumsden A, Nik SHM, et al. Accelerated brain aging towards transcriptional inversion in a zebrafish model of the K115fs mutation of human PSEN2. PLOS ONE. 2020;15(1):e0227258.

63. Wickham H. ggplot2: Elegant Graphics for Data Analysis. Springer-Verlag New York; 2016.

64. Kolde R. pheatmap: Pretty Heatmaps. 1.0.12 ed 2019.

65. Conway JR, Lex A, Gehlenborg N. UpSetR: an R package for the visualization of intersecting sets and their properties. Bioinformatics. 2017;33(18):2938–40.

